# Motif Interactions Affect Post-Hoc Interpretability of Genomic Convolutional Neural Networks

**DOI:** 10.1101/2024.02.15.580353

**Authors:** Marta S. Lemanczyk, Jakub M. Bartoszewicz, Bernhard Y. Renard

**Affiliations:** Hasso Plattner Institute for Digital Engineering, Digital Engineering Faculty, University of Potsdam; Computer Science and Artificial Intelligence Laboratory, Massachusetts Institute of Technology

## Abstract

Post-hoc interpretability methods are commonly used to understand decisions of genomic deep learning models and reveal new biological insights. However, interactions between sequence regions (e.g. regulatory elements) impact the learning process as well as interpretability methods that are sensitive to dependencies between features. Since deep learning models learn correlations between the data and output that do not necessarily represent a causal relationship, it is difficult to say how well interacting motif sets are fully captured. Here, we investigate how genomic motif interactions influence model learning and interpretability methods by formalizing possible scenarios where interaction effects appear. This includes the choice of negative data and non-additive effects on the outcome. We generate synthetic data containing interactions for those scenarios and evaluate how they affect the performance of motif detection. We show that post-hoc interpretability methods can miss motifs if interactions are present depending on how negative data is defined. Furthermore, we observe differences in interpretability between additive and non-additive effects as well as between post-hoc interpretability methods.

## Background

Convolutional neural networks (CNN) excel at various sequence-based tasks due to their capability to learn patterns and complex interactions making these models an efficient method for many predictive tasks in the field of genomics [1]. However, for many biological applications, predictions alone are insufficient for understanding the underlying mechanisms for a given problem [2]. Besides verifying that a model learned meaningful predictions, interpreting CNNs can lead to new insights for genomic questions [3, 4]. With the help of such interpretations, the models’ decisions can be verified, or new insights can be obtained [5]. To explain the outcome of a CNN, post-hoc interpretation methods are a commonly used approach. Instead of training intrinsically interpretable models, post-hoc interpretation methods are applied after the training process on a fully trained model. Known methods include feature permutation [6], Integrated Gradients [7], and DeepLIFT [8]. When applied to sequence data, scores are assigned to each position in an input sequence based on their contribution to an individual prediction. By aggregating attribution scores from multiple input sequences, it is possible to extract meaningful motifs [9]. Identified motifs can be compared with known motifs in task-specific databases to determine their biological relevance by using methods for motif comparison like TOMTOM [10, 11].

The quality of contribution scores can be affected by multiple factors besides model complexity so the evaluation of interpretability performance for machine learning models is crucial [12, 13, 14]. This also applies to interpretability for biological neural networks. Some differences in contribution scores can be attributed to the architectural choices for the model. In [15] it was shown that exponential activation functions can lead to more interpretable motif representations in first-layer filters than for other functions, like ReLU, sigmoid or tangent activation function, as well as specific choices for filter size, max pooling width and model depth [16]. Interpretability can be also improved by introducing robustness with the help of regularization, random noise injection, and adversarial training [17]. While those approaches improve interpretability in general, one has to keep in mind that post-hoc interpretations represent what a model learned. The learned correlative features do not necessarily represent causal effects. Dependencies between features complicate learning causal effects with machine learning models. Not only are many features in biological sequences in relationship with others, but groups of such locally dependent features can also be part of a higher interaction representing a regulatory logic hidden in deeper layers [18]. Such biological interaction can be, for instance, cooperative binding of transcription factors to DNA [19]. This can result in misleading explanations if the complete underlying biological mechanism is not uncovered by the model. Furthermore, some of the attribution methods are based on the assumption that input features are independent. Dependencies between motifs can influence attribution methods so that subsets of interacting motifs can produce incomplete or noisy interpretations. It is crucial to analyze if post-hoc methods are capable of capturing interacting motifs and, therefore, the underlying causal effects in an understandable manner.

To tackle those challenges, we design suitable data for different sources of interactions and evaluate the interpretability performance of models trained with that data. For that, we first investigate how interactions affect model training and interpretability in general by using different negative data sets, forcing the model to learn interactions explicitly or not (Fig. 1, left). The selection of negative data for machine learning is an ongoing problem for various biological tasks [20, 21] since it can influence the discriminative power of models. One mistake is not to include data where there is some uncertainty for the data label, for example, due to the similarity between positive and negative instances. Using easily distinguishable data results in a simpler training task and can still lead to good predictions if the new data is similarly structured to training data. However, the predictive performance decreases when the model is confronted with uncertain data like novel populations. In the case of genomic sequences, this can happen when using random sequences for the negative training data set instead of carefully curated samples. Models that are trained on non-random negative data have to learn more complex relationships between motifs in the positive class in contrast to random negative sequences, which allow the model only to learn subsets or even individual motifs to distinguish between classes. While the influence of negative data on prediction tasks is a known problem, it is not well explored how it influences interpretability.

**Figure 1.**
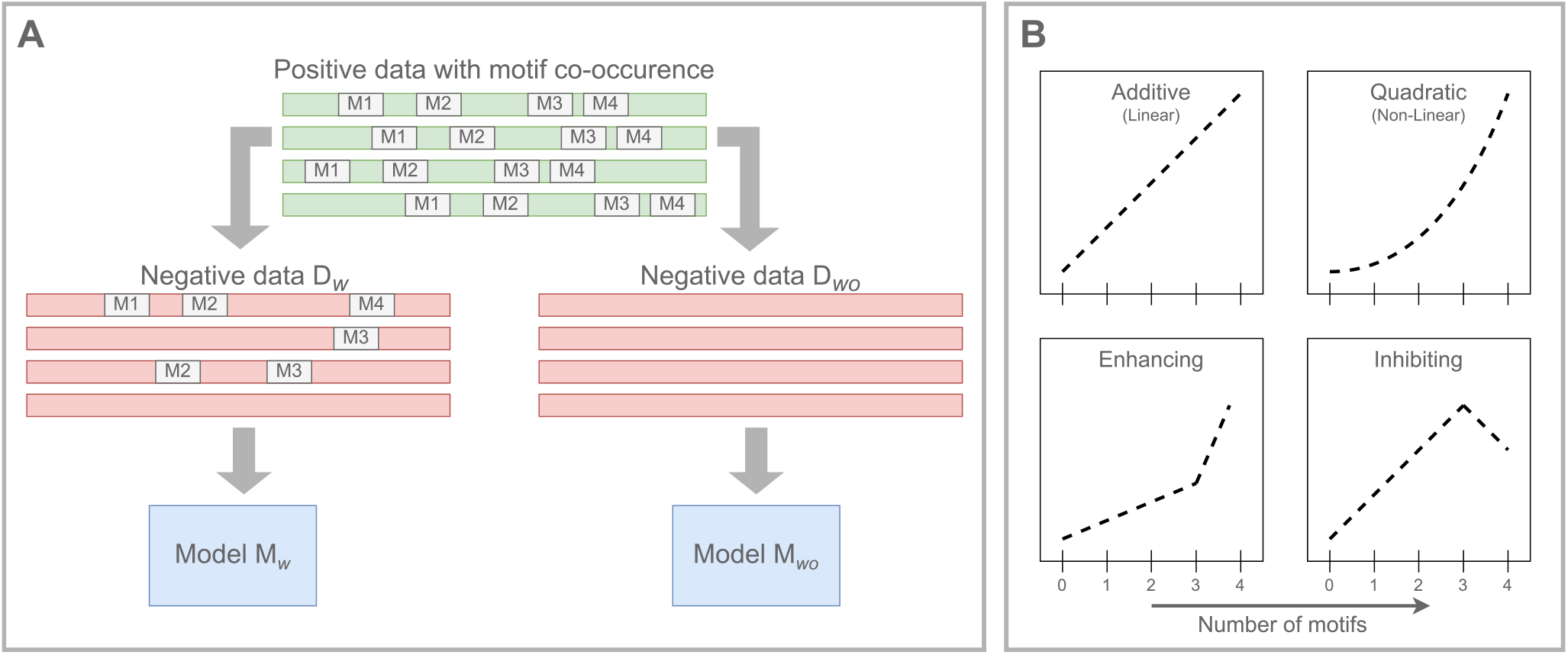
Data design for motif interactions. **(Classification)** Capturing a motif interaction in the form of co-occurrence does not only depend on the positive dataset containing all motifs. Depending on the nature of the negative data, models differ in how the class boundaries and the interactions are learned. Here, we also analyze the impact on interpretability. **(Regression)** Deep learning models are capable of learning interactions with varying complexities. We investigate if complexity also influences post-hoc interpretability. Detailed experiment and data generation settings can be found in Fig. 4.

Secondly, we look at the interaction effects between motifs (Fig. 1, right). In regulatory genomics, different experiments allow us to determine the functions and characteristics of non-coding DNA regions [22]. This includes, among other methods ChIP-seq, DNase-seq, and ATAC-seq where the enrichment of sequence fragments is measured for various functions like protein binding locations or chromatin accessibility. The output is then mapped to the genome resulting in per-position counts, which can then be binarized based on signal peaks representing the sites of interest. Previous deep learning models focused on the prediction of binarized peaks [23] and perform, therefore, a classification problem. Currently, many state-of-the-art models are used to directly predict the signal of an assay rather than just the presence of a peak so that the output gives a more precise prediction of the signal of interest [24]. However, the regression task is more complex since the model needs to learn a function between the input sequence and the numerical output instead of just distinguishing between classes. Since multiple regulatory elements can be involved in a regulatory mechanism, interactions between motifs complicate the prediction task. Motif interactions can occur in multiple forms, including additive effects as well as multiplicative interactions [25]. Here, we explore if the complexity of interaction effects influences model interpretability.

## Results

We evaluated the influence of motif interactions on motif detection by first defining various interactions and then simulating data containing those interactions. Model architecture and training setup are described in the CNN section. We evaluated the model performance with regard to prediction and motif detection capability as well as the post-hoc attribution methods with metrics introduced in the evaluation section.

The motif set used for the following evaluation consists of motifs in table 1 obtained from the JASPAR database for transcription factor binding sites [26].

**Table 1.** Motif data sets. The data set *‘distinct_1’* was used for the evaluation of heterologous motifs while the dataset *‘MEF2A’* represents homologous data sets.

| distinct_1 |  | MEF2A |  |
| --- | --- | --- | --- |
| Motif name | Motif ID | Motif name | Motif ID |
| CREB1 | MA0018.3 |  | MA0052.1 |
| NHLH1 | MA0048.1 | MEF2A | MA0052.2 |
| ETS1 | MA0098.3 |  | MA0052.3 |
| TEAD1 | MA0090.3 |  | MA0052.4 |

### Negative Sequences

To evaluate the influence of negative data on model interpretability, we simulate the scenario in a classification problem predicting an outcome based on the co-occurrence of a set of motifs. For that, we create two data sets that contain the same sequences for the positive class but differ in their negative sequences. Specifications can be found in the respective method section. The positive data set is described by the co-occurrence of all *n* motifs that are here set to *n* = 4. Regarding the negative data, we distinguish between the data set containing 0 to *n−* 1 motifs and a data set containing only random sequences without any motifs inserted. We call the model trained on data with motif subsets in the negative data set *ℳ*_*w*_ and the model without motifs *ℳ*_*wo*_.

Based on the negative data set used for training, models learn different ways to predict the positive class. To investigate the underlying learning mechanisms, we use negative test data similar to the training data set with motif subsets in the negative data set. Each possible combination of motifs is represented by the same number of sequences in the negative data set, as can be seen in Fig. 2A. The respective accuracy can be seen in Fig. 2B. While both models have a decent accuracy for the positive data (*acc*(*ℳ*_*w*_) = 0.9485, *acc*(*ℳ*_*wo*_) = 0.9992), the models differ in the accuracy for the negative data set (*acc*(*ℳ*_*w*_) = 0.9112, *acc*(*ℳ*_*wo*_) = 0.5916, more detailed accuracies can be found in Table 2).

**Table 2.** Recall for CNNs based on data with negative sequences containing motif subsets (*M*_*w*_) and without motif subsets (*M*_*wo*_). While *M*_*w*_ is capable of distinguishing between positive sequences (4 motifs) and negative sequences (0-3), *M*_*wo*_ has low accuracies for sequences with 2 or 3 motifs since sequences with subsets of the interacting motif set were not present during training. The number of filters has no large influence on the accuracy.

| (a) MEF2A |  |  |  |  | (b) Distinct 1 |  |  |  |  |
| --- | --- | --- | --- | --- | --- | --- | --- | --- | --- |
| MEF2A | large (32 filter) |  | small (4 filter) |  | distinct_1 | large (32 filter) |  | small (4 filter) |  |
| #Motifs | $M_w$ | $M_{wo}$ | $M_w$ | $M_{wo}$ | #Motifs | $M_w$ | $M_{wo}$ | $M_w$ | $M_{wo}$ |
| 4 | 0.9832 | 0.9992 | 0.9772 | 0.9999 | 4 | 0.9307 | 0.9994 | 0.9485 | 0.9992 |
| 3 | 0.91 | 0.034 | 0.8965 | 0.04 | 3 | 0.6815 | 0.0615 | 0.7215 | 0.089 |
| 2 | 0.9973 | 0.3336 | 0.9963 | 0.4534 | 2 | 0.9893 | 0.6177 | 0.9647 | 0.6 |
| 1 | 1 | 0.91 | 1 | 0.961 | 1 | 1 | 0.988 | 0.9985 | 0.9795 |
| 0 | 1 | 1 | 1 | 1 | 0 | 1 | 0.998 | 1 | 1 |

**Table 3.**
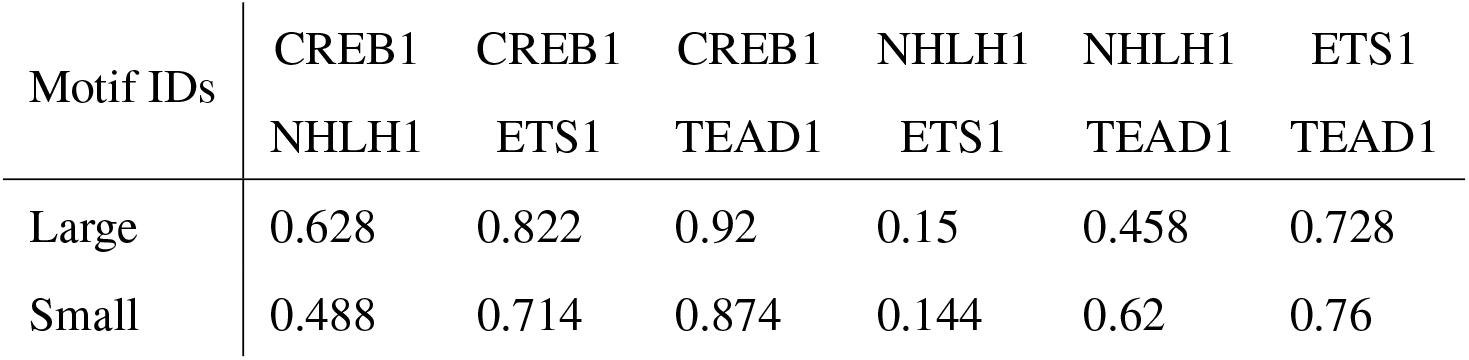
Subset recall for *M*_*wo*_ trained on the *distinct_1* data set. Each test sequence contains 2 motifs. The accuracies are calculated for each pairwise motif combination to see if the subsets meet the overall accuracy (Large model: 0.6177 and Small: 0.6) or if there are preferences in motifs. Low accuracy means that many sequences were predicted as positive and, therefore, that motif subset is evidence for the positive class for the model.

**Table 4.** Motif-wise AUPRC values for contribution scores for models trained on MEF2A homologous data set. The scores are shown for positive sequences containing all 4 interactive motifs.

|  |  | MA0052.1 | MA0052.2 | MA0052.3 | MA0052.4 |
| --- | --- | --- | --- | --- | --- |
| DeepLIFT | w | 1.0 | 0.7445 | 0.9006 | 0.7073 |
|  | wo | 1.0 | 0.8085 | 0.9743 | 0.7429 |
| Feature Permutation | w | 0.9513 | 0.6713 | 0.7903 | 0.5942 |
|  | wo | 0.9463 | 0.7787 | 0.9049 | 0.7165 |
| Integrated Gradients | w | 1.0 | 0.7268 | 0.8819 | 0.6698 |
|  | wo | 1.0 | 0.8317 | 0.9933 | 0.7741 |

**Figure 2.**
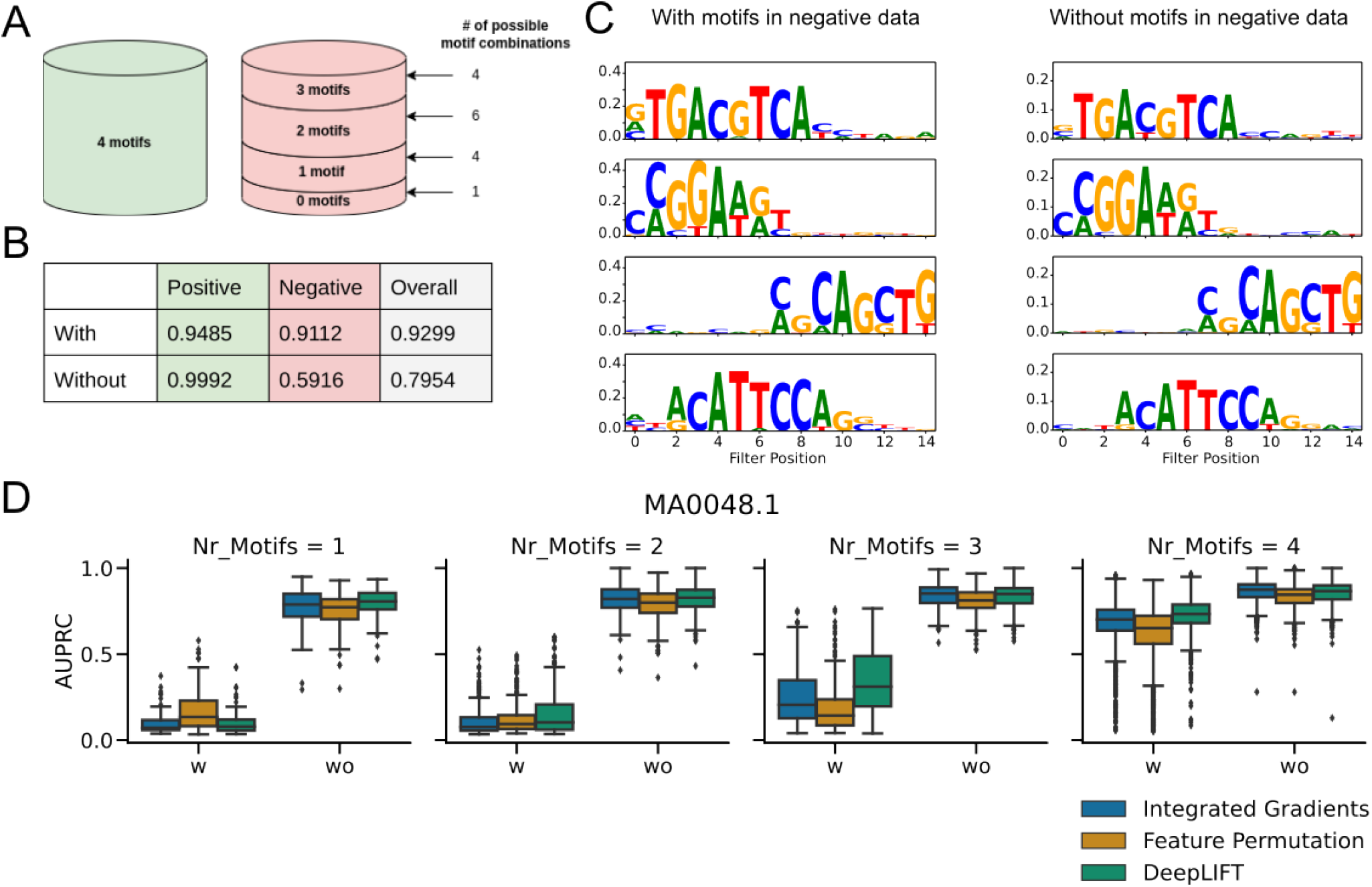
Results for models trained on two different negative sequence data sets to predict the co-occurrence of 4 motifs. **A** Both model types are trained on the same positive data containing sequences with 4 motifs inserted. One model type (*M*_*w*_) additionally includes motif subsets with max. 3 motifs in the negative data set as depicted on the right side, while the second only includes random sequences without any motifs (*M*_*wo*_). **B** Predictive accuracy for both models. While both models have good accuracy for the positive class containing 4 motifs, the model *M*_*wo*_ performs poorly on the negative data, as expected. **C** Weights of convolutional layer filters. Both models learned similar representations of the motifs within the layer. **D** AUPRC values for contribution scores for positive class. High values indicate high contribution scores for motif positions compared to random sequences and, therefore, better motif detection. For *M*_*w*_, the AUPRC scores decrease significantly for models containing only subsets of motifs which indicates a lower motif detection capability.

We trained models with a minimal number of filters so that the number of filters equals the number of motifs (#filters = 4), as well as CNNs with 32 filters. Both minimal filter models captured nearly identical motif weights in the convolutional filter (Fig. 2C), showing that both models learned a similar representation of the inserted motifs.

We calculated the contribution scores for test sequences using DeepLIFT (DL), Integrated Gradients (IG), and Feature Permutation (FP). Since the ground truth is known for our simulated data, we can quantify how well the contribution scores were assigned to the motifs. The AUPRC value (see methods) should be high if an attribution method assigned high absolute scores to the motif positions compared to random positions. By using the absolute scores, the negative influence of the motifs is also captured.

In Fig. 2D, AUPRC scores are shown for the dataset *distinct_1* for the NHLH1 Motif (ID: MA0048.1). The test sequences contained subsets of motifs, of which at least one was the motif of interest. For the positive data with all motifs present (2D, iv), differences can be observed between the AUPRC medians of both models with *M*_*w*_ having lower AUPRC (Δ*IG* : 0.173931, Δ*DL* : 0.194141, Δ*FP* : 0.133069, see Table 5). The motif detection performance for *M*_*w*_ drops for the negative sequences containing 1 to 3 motifs, while the performance for *M*_*wo*_ remains more stable for all sequences (2D, i-iii). There are no major differences between the attribution methods for motif NHLH1 (see 5). Similar observations can be made for the other motifs (see Supplement Fig. 5). Besides the data sets containing heterologous motifs, data sets with homologous motifs were investigated. Similar to the models based on heterologous motif sets, motif detection performance decreases for *M*_*w*_ the fewer motifs are present in the sequence. However, for all motifs (MEF2A), an increase in the AUPRC scores can be observed for Feature Permutation when 1 or 2 motifs are present while the performance for *M*_*wo*_ drops.

**Table 5.** AUPRC values for contribution scores for models trained on *distinct_1* heterologous data set for one motif (NHLH1). *M*_*w*_ has lower values, especially for negative data containing 1-3 motifs.

| #Motifs | Integrated Gradients |  | Feature Permutation |  | DeepLIFT |  |
| --- | --- | --- | --- | --- | --- | --- |
| | $M_w$ | $M_{wo}$ | $M_w$ | $M_{wo}$ | $M_w$ | $M_{wo}$ |
| 4 | 0.701 | 0.8749 | 0.6513 | 0.8454 | 0.7331 | 0.8661 |
| 3 | 0.2053 | 0.8529 | 0.1432 | 0.8124 | 0.3106 | 0.85 |
| 2 | 0.0765 | 0.8213 | 0.094 | 0.7998 | 0.1032 | 0.8279 |
| 1 | 0.0720 | 0.7881 | 0.1346 | 0.772 | 0.0789 | 0.8061 |

### Interaction effects

As described in the method section, various interaction effects based on the co-occurrence of an interacting motif set are represented by a function that generates labels for the input sequences. For that, we assigned each sequence a numerical label based on the number of motifs contained in the input sequence. We distinguish between an additive, enhancing, quadratic, and inhibiting effect. Further details can be seen in Fig.1 (right). To make the prediction accuracy comparable, we bin the outcomes around the possible labels and calculate if the predicted value falls in the interval. We averaged the accuracy and the recall as well as the AUPRC across 5 models with different seeds to decrease potential noise. We compare the AUPRC values for all effects individually for each motif. We also differentiate between a model with 4 and 32 filters.

We calculate the recall separately for each subset depending on the number of containing motifs (see Table 6). The total accuracy for each effect lies between 0.8599 and 0.8729 for the model with 32 filters. However, there are differences between the effects when it comes to recall. The additive interaction effect has a mostly stable performance between subsets, while the other effects vary in recall. Additionally, the performance for negative sequences without any motifs is low for all effects ranging between 0.3588 to 0.6568.

**Table 6.**
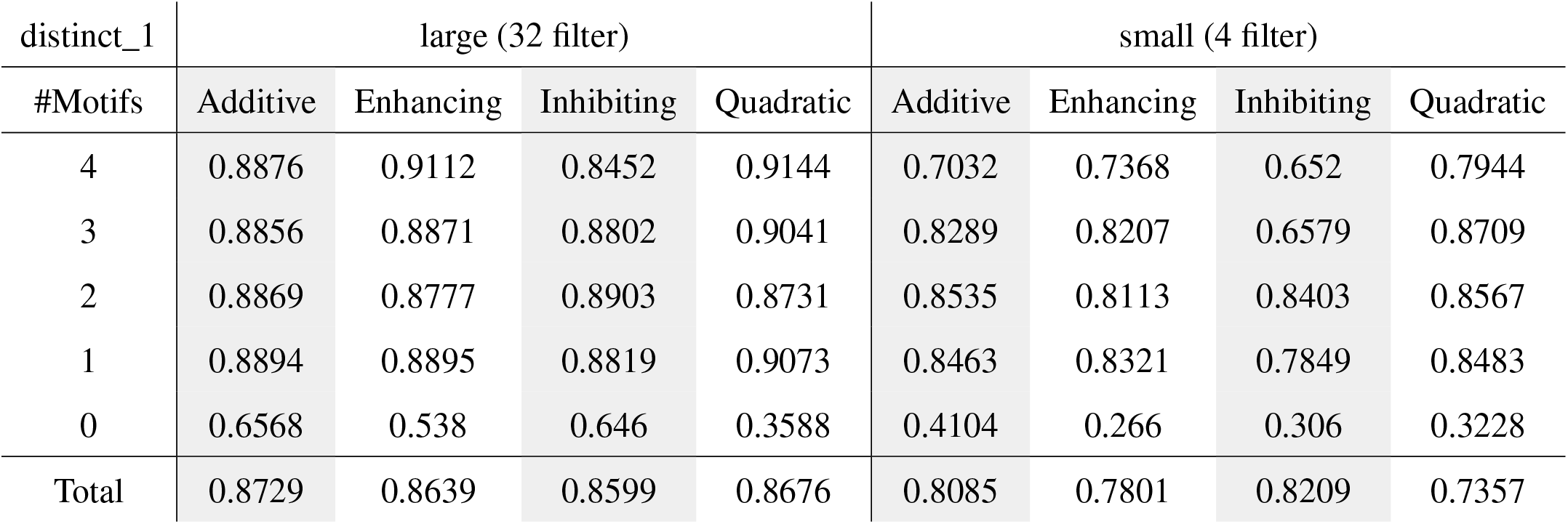
Subset recall and total accuracy for regression models trained on *distinct_1*.

| distinct_1 | large (32 filter) |  |  |  | small (4 filter) |  |  |  |
| --- | --- | --- | --- | --- | --- | --- | --- | --- |
| #Motifs | Additive | Enhancing | Inhibiting | Quadratic | Additive | Enhancing | Inhibiting | Quadratic |
| 4 | 0.8876 | 0.9112 | 0.8452 | 0.9144 | 0.7032 | 0.7368 | 0.652 | 0.7944 |
| 3 | 0.8856 | 0.8871 | 0.8802 | 0.9041 | 0.8289 | 0.8207 | 0.6579 | 0.8709 |
| 2 | 0.8869 | 0.8777 | 0.8903 | 0.8731 | 0.8535 | 0.8113 | 0.8403 | 0.8567 |
| 1 | 0.8894 | 0.8895 | 0.8819 | 0.9073 | 0.8463 | 0.8321 | 0.7849 | 0.8483 |
| 0 | 0.6568 | 0.538 | 0.646 | 0.3588 | 0.4104 | 0.266 | 0.306 | 0.3228 |
| Total | 0.8729 | 0.8639 | 0.8599 | 0.8676 | 0.8085 | 0.7801 | 0.8209 | 0.7357 |

**Table 7.** Subset recall and total accuracy for regression models trained on *MEF2A*.

| distinct_1 | large (32 filter) |  |  |  | small (4 filter) |  |  |  |
| --- | --- | --- | --- | --- | --- | --- | --- | --- |
| #Motifs | Additive | Enhancing | Inhibiting | Quadratic | Additive | Enhancing | Inhibiting | Quadratic |
| 4 | 0.9492 | 0.9576 | 0.9416 | 0.9576 | 0.9284 | 0.9328 | 0.9196 | 0.9256 |
| 3 | 0.9401 | 0.9457 | 0.9341 | 0.9458 | 0.9128 | 0.8956 | 0.9024 | 0.9036 |
| 2 | 0.9361 | 0.9245 | 0.9437 | 0.923 | 0.8992 | 0.877 | 0.9049 | 0.8832 |
| 1 | 0.9086 | 0.9105 | 0.909 | 0.9243 | 0.8795 | 0.8723 | 0.8688 | 0.8974 |
| 0 | 0.8612 | 0.8504 | 0.8588 | 0.74 | 0.8156 | 0.7656 | 0.8144 | 0.6484 |
| Total | 0.9190 | 0.9177 | 0.9174 | 0.8981 | 0.8871 | 0.8687 | 0.88201 | 0.8516 |

AUPRC scores are compared between the interaction effects as well as interpretability methods and motifs. For IG, The motif-wise AUPRC for the additive effect is high for all motifs with median values between 0.86 and 1, similarly for the inhibiting effect (0.85-0.98). For the enhancing and quadratic effect, the AUPRC values do not show a large decrease for motifs NHLH1 and ETS1, whereas for motifs CREB1 and TEAD1 the values drop, especially for the quadratic interaction (AUPRC median: 0.75 for CREB1 and 0.68 for TEAD1) including an increased variance in AUPRC scores. Feature Permutation performs in total worse than Integrated Gradients and DeepLIFT having a large variance with median values around 0.5 to 0.85. Here, the highest values were assigned to the inhibiting effect while Feature Permutation also performed worse for other non-additive effects. DeepLIFT shows the most robust AUPRC along the motifs and effects. However, the assigned values to motifs CREB1 and TEAD1 are slightly worse for the non-additive effects, including the inhibiting effect.

### Subsets of interactive motifs in negative data can lead to different decision boundaries and therefore to varying motif detection performances

While the performance for positive sequences with all motifs from the interactive set is similar for both models, there are strong differences for the negative sequences when it comes to accuracy and motif detection performance. As expected, *M*_*wo*_ is not capable of classifying all negative sequences containing subsets of motifs correctly since there were no sequences with subsets in the training data. Accuracy drops with increasing size of the motif subset. The accuracy for sequences containing only 1 motif is still high (*acc*_1 motif_ = 0.9795), while for sequences with 2 and 3 motifs, the model performs poorly (*acc*_2 motifs_ = 0.6, *acc*_3 motifs_ = 0.089) compared to *M*_*w*_ (*acc*_2 motifs_ = 0.9647, *acc*_3 motifs_ = 0.7215). This indicates that the model *M*_*wo*_ does not generalize well and learns only subsets of the interactive motif sets to classify a sequence as positive instead of the full motif set like *M*_*w*_. The accuracy scores for sequences containing 2 motifs differ depending on the present motif subset. For instance, the averaged accuracy for the motif set containing motifs CREB1 and TEAD1 reaches 0.874 while for the set containing NHLH1 and ETS1 only 0.144 (see all accuracies in 3). Since the negative sequences containing 3 motifs are mostly classified as positive, and most sequences containing 1 motif are correctly classified as negative, those observations suggest that the decision boundary between classes is based on specific motif sets containing between 2 and 3 motifs.

As we can see in the accuracy values, *M*_*w*_ has to learn a decision boundary between sequences containing 3 and 4 motifs. If we permute one motif (which equals a motif removal) in the positive class sequence, the outcome should change to negative. Here, the permuted motif has therefore a high contribution to the outcome. However, if we look at a negative sequence containing 3 motifs and permute one motif, the resulting sequence with 2 motifs still belongs to the negative class. The permuted motif does not contribute to an outcome change which can result in low AUPRC scores. For *M*_*wo*_, it is unclear how the decision boundary is learned and which individual motifs or motif subsets must be present to influence the model’s decision. Based on the observations in the accuracies, the presence or absence of individual motifs could already impact the outcome in sequences with fewer motifs. That impact could be reflected in the higher AUPRC scores also for the negative sequences.

### Non-additive interactions can influence interpretability independent of accuracy

We observe a decrease in AUPRC values for non-additive interaction effects for the models with 32 convolutional filters (see Fig. 3 A). DeepLIFT performed the most stable when it comes to interpretability, with only a small decrease between the additive effect and the non-additive ones. On the other hand, IG performed worse on motifs CREB1 and TEAD1 for the enhancing and the quadratic effect, while there is no large difference for motifs NHLH1 and ETS1 between the different interaction effects. Feature Permutation has overall low AUPRC values suggesting that it is not suitable for the regression task. We also included the absolute errors of the predictions to validate if lower AUPRC values result from bad predictions. The absolute errors of the predictions for the sequences with low AUPRC values for the quadratic effect do not show an increase (see Fig. 3 B). Therefore, we can assume that the worse performance in detecting the motifs can result from the more complex interaction independent of the accuracy.

**Figure 3.**
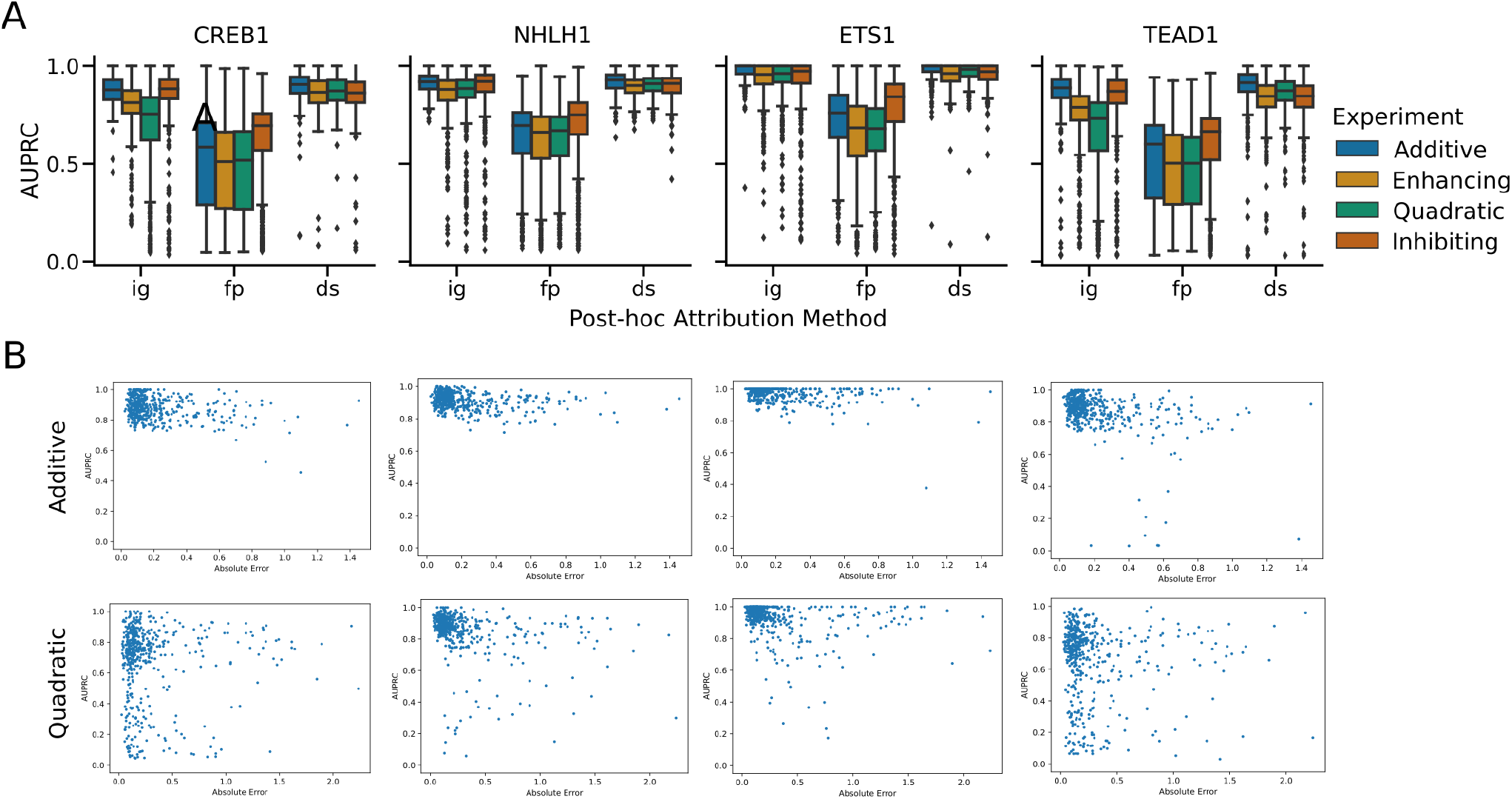
Results for interaction effects for sequences containing all 4 motifs. **A** Motif-wise AUPRC values for heterologous motif set. For Integrated Gradients a drop in AUPRC values can be observed for the enhancing and quadratic effect for motifs for CREB1 and TEAD1 compared to motifs NHLH1 and ETS1. Only a small decrease can be observed for the non-additive interactions for DeepLIFT. Feature Permutation performs overall worse for the regression task having high variance in the scores. **B** Comparison between Accuracy and AUPRC values for the additive and quadratic effect for Integrated Gradients. Low AUPRC values for the quadratic effect do not show increased error values.

## Discussion

Interpretation has become a crucial part of deep learning applied in the field of genomics. While the validation of identified motifs can confirm that a model learned meaningful patterns, the lack of complete biological data makes it difficult to prove the completeness of all motifs and therefore derive causal insights. Here we investigated from a data-centric point of view how interactions in genomic datasets can result in missing or noisy interpretations.

We concentrate in this work on the effects of the co-occurrence of motifs and their effects on the outcome. However, there are further aspects that can influence motif interactions. For example, the cooperativity of transcription factor motifs can additionally depend on order, orientation, and distance (including 3-D genome distances) between regulatory elements [27] as well as temporal causes [28]. Motifs in our generated data sets have a fixed order and orientation as well as the same distance between each other for all sequences in one data set. In this way, we focus solely on a fixed grammar to reduce complexity and other factors that could affect interpretability. It is also important to point out that the model might see the tasks more as a classification counting the number of motifs instead of learning a function since the labels are discrete based on the number of motifs in the sequence. In this case, the kind of interaction function might not be relevant to the model since it is not learning the function itself. Including other information like distances between motifs could improve the simulation by making a long function continuous and, therefore, additive and non-additive interaction effects could be better explored.

In the negative sequence experiment, we observe a trade-off between accuracy and motif detection, especially for negative data. Here, multiple data augmentation strategies can be evaluated to obtain better motif detection performance or desired interpretability outcomes on interactive data while preserving biological functionality (e.g. [29]). Motif detection can also be improved by accounting for interactions like in [18] where stochastic masks are used to find sets of motif features that preserve or change the outcome and therefore avoid saturation effects.

In this work, we focused on simple CNNs to break down the problem of interpretability to interactions. Using more complex architectures would result in additional sources affecting interpretability so it would be more difficult to separate the interaction effects from the other sources. However, currently, more complex models with more sophisticated modules such as the attention mechanism are applied to genomic problems to capture interactions within genomic sequence data (see an overview on genomic large language models (LLM) [30]). So far, many approaches to interpreting genomic LLM models focus on the analysis of the attention scores or the output with post-hoc methods that mostly offer interpretations on the input token level. One ongoing challenge is to uncover the grammar between interacting motifs so that interpreting genomic LLMs beyond those approaches could give better explanations of underlying biological processes. Also, pre-training of genomic LLMs should be explored in the context of interactions. Especially, if downstream tasks are missing relevant data, like in the negative data experiment, it is necessary to analyze how the missing information is imputed.

As we could also see in our results, machine learning models do not necessarily learn the underlying causal effect of biological mechanisms. Thus interpreting models after training is not always suitable for knowledge extraction. Therefore, designing interpretable architectures that capture the interactions explicitly instead of only relying on post-hoc model interpretation could be a better approach for motif identification as well as interaction detection.

## Conclusion

We analyze the influence of motif interactions on post-hoc interpretability methods. First, we investigate how motif co-operativity can affect model learning depending on how interacting motifs are present in the negative data set. We observe that interpretability performance can decrease when interactions are learned more explicitly by the model. Especially for negative sequences, evidence for the positive class can be missed. Secondly, we formalize different interaction effects (additive as non-additive) and compare those with regard to interpretability. We discovered differences between the effects as well as the interpretability methods, from which we deduce that post-hoc interpretability is affected by complex interactions.

## Methods

### Motif interactions

We define each prediction task as a function *F* : *X → Y*. The input *x ∈ X* = *{*0, 1*}*^*n*^ describes the presence or absence for all motifs *i ∈ M* = *{*1,…, *n}* in the input sequence. If a motif *i* is present in a given sequence, then *x*_*i*_ = 1, if absent then *x*_*i*_ = 0. The outcome 𝕐 depends on the task. For regression problems, we define *Y* = ℝ, while *Y* = *{*0, 1*}* applies for binary classification tasks.

### Interaction effect on outcome

Motif interactions, eg. co-occurrence, can be expressed as logical constructs using `AND, OR`, and `NOT`. The nature and magnitude of the effects of these relationships on the outcome (eg. non-linearity, inhibition, activation) can be encoded in the target values of a regression task. Since we assume that there are no other features that can influence the outcome except the given motifs, we derive the following definitions from the definitions in [31].

Let *F*(*x*) be the sum of the effects of all possible subsets of motifs, where each motif combination has its influence on the outcome:

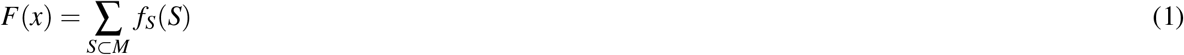

The independent effect of a motif *m*_*i*_ on *F*(*x*), which does not rely on the presence or absence of other motifs, is called the main effect and is defined here as a subfunction *f*_*i*_(*x*_*i*_). If *F*(*x*) is only affected by the main effects of the motifs and therefore does not contain any interactions between them, the function is described as an additive interaction:

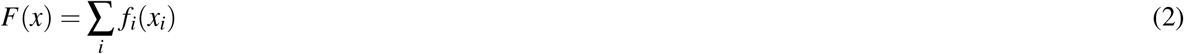

The task contains at least one non-additive interaction if there is a subset *S ⊂ M*, |*S*| *≠*1 so that *f*_*S*_(*S*) *≠*0 and therefore *F*(*x*)*≠*∑_*i*_ *f*_*i*_(*x*_*i*_) [32]. In that way, we can define different interactions as functions. We show two examples that we use for our evaluation.

#### Example 1: Enhancement and Inhibition

Besides the main effects of individual motifs, we introduce an enhancement/inhibition term so that

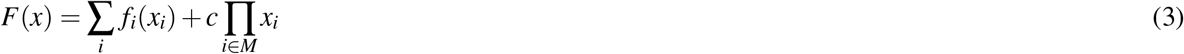

for some constant *c ∈* ℝ *\ {*0*}*. Here, a non-zero value is added to the outcome if and only if the input sequence contains all motifs from the interacting motif set. The co-occurrence of all motifs in that set enhances the individual main effects on the outcome.

#### Example 2: Non-linear relationship

The relationship between motifs can also be expressed with a non-linear function depending on the subsets of the interacting motif set. As an example of a nonlinear interaction, we use a quadratic relationship:

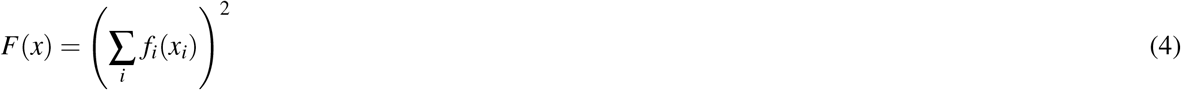

Combinations of different interactions are also possible and add complexity to the task. However, we use the described interactions to investigate the differences between additive and non-additive interactions.

### Negative sequences

The performance of a model strongly depends on the available data. The nature of the data set can have different effects on how the model learns the interactions between motifs. Here, we simulate a binary classification problem with positive and negative classes. While the data for the positive class plays a major role in binary classification, the negative class can also impact the resulting model. In this case, the positive class represents the co-occurrence of all motifs in the interacting motif set *M* so that *∀i ∈ M* : *x*_*i*_ = 1. Different negative data sets can be chosen for the same task while the model can still have a similar predictive performance in the end. We distinguish between two different negative data sets. One data set includes individual motifs or subsets of the interacting motif set in the negative data set so that *∃i ∈ M* : *x*_*i*_ = 0. In contrast, sequences from the second negative data set do not contain any motifs from the interacting motifs set and therefore *∀i ∈ M* : *x*_*i*_ = 0.

### Sequence data

Genomic sequences have to be transformed into numerical matrices so they can be processed by CNNs. Each column of this matrix stands for one sequence position where the base at this position is represented by a one-hot-encoding vector. We use sequences with the length of 250 base pairs resulting in matrices with the size of 4x250.

We obtain real transcription factor binding motifs from the JASPAR database [26] for the evaluation. We distinguish here between subsets of homologous and heterologous motif subsets to investigate if motif similarity influences interpretability. The similarity was measured by the Pearson correlation coefficient for motif similarity [10]. We used the implementation from the biopython package [33]. Motifs can have different lengths, e.g. transcription factor binding sites have a length of around 5-31 nt [34], which may also influence interpretability if cooperating motifs differ in size. We picked for our experiment motifs with approximately similar lengths. The selected motifs can be found in Table 1. For each task, a grammar is generated with a fixed order of motifs and distances between the motifs. The grammar itself is inserted randomly within the sequence to ensure invariance regarding the position of the motif grammars. The distance between the motifs is larger than the filter size, so the filters learn individual motifs, not overlapping regions. Labels were generated by following the interaction definitions above. Training, test, and validation sequences are the same for models that are compared, and the data sets only differ in the labels that encode the interactions.

### Convolutional Neural Networks

To ensure that differences in interpretability performance cannot be traced back to differences in predictive performance, one requirement is that models that are compared have similar performance.

We use CNNs with one convolutional layer with filters approximating the length of the chosen motifs to learn localist representations as described in [16]. 3 dense layers follow the convolutional layer to learn the interactions between the motifs. We apply batch normalization on the inputs before passing them to a ReLU function. Additionally, we apply max pooling in the convolutional layers.

### Interpretability Methods

We use feature permutation (FP) [6], Integrated Gradients (IG) [7], and DeepLIFT (DL) [8] as post-hoc attribution methods. The analyses are performed with the method implementations from the Captum library for PyTorch [35]. Since we obtained similar results for average and zero reference sequences, we use the zero reference sequence due to the shorter computational time. Contribution scores are averaged over the 5 models to reduce noise.

## Evaluation

Contribution scores were evaluated similarly to [15]. Each position in the sequence gets a label assigned depending on if it belongs to a motif (1) or not (0). Contribution scores of motif positions are then compared to those of random positions by calculating the AUPRC (see overview of the model evaluation in Supplement Fig. 4). AUPRC scores are calculated for each motif separately. AUPRC scores were then visualized via boxplots. Interpretability performance was then compared between the models of interest by analyzing the differences of the AUPRC on the same sequence test set.

**Figure 4.**
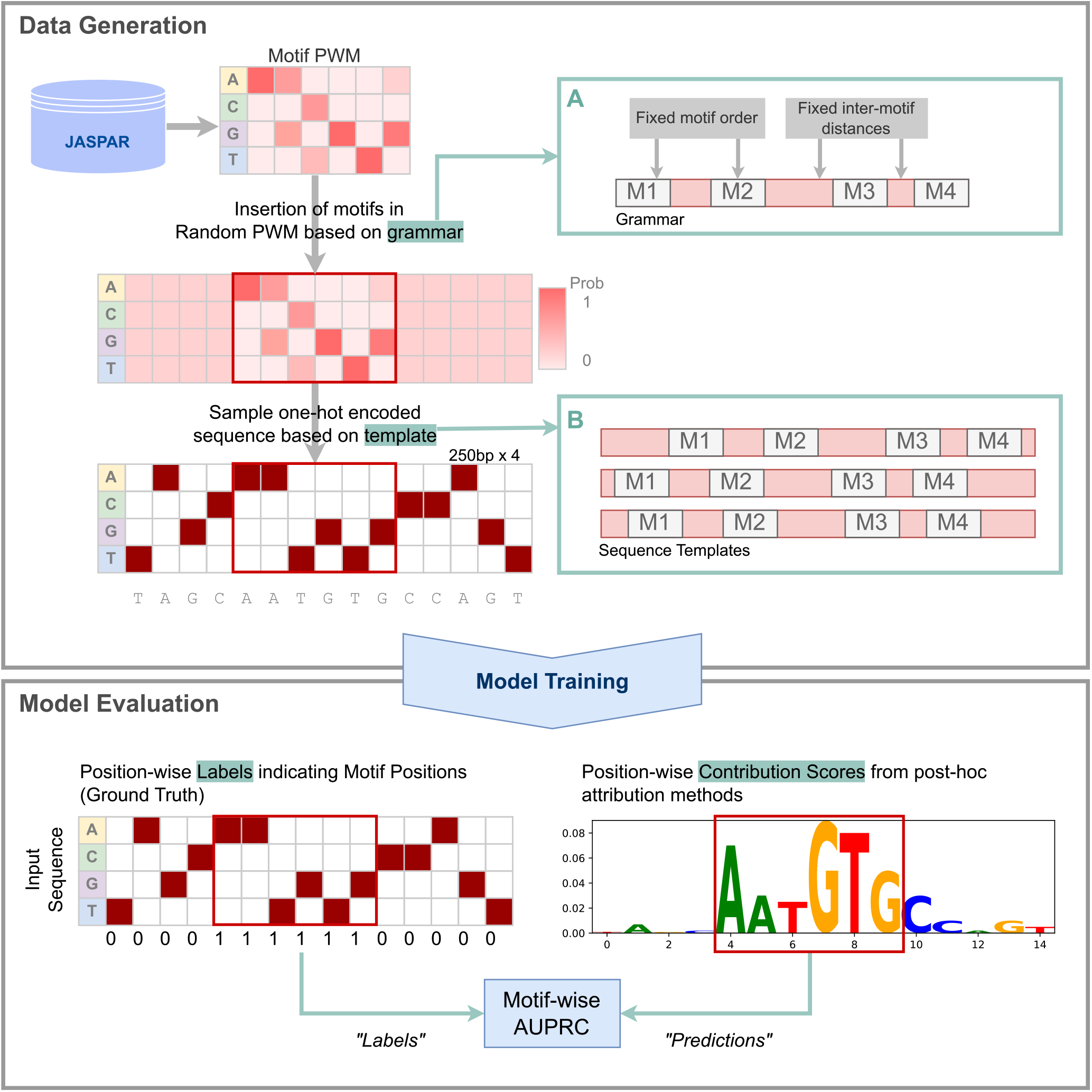
**Data Generation** We obtained the PWMs for 5 motif sets (see Table 1) from the JASPAR database. We create grammars for each set which consist of a specific motif order and distances between motifs (A). The presence of a motif depends on the investigated interaction (see methods section). One-hot-encoded sequence templates in the form of PWMs are generated for each input sequence from the grammar (B) from which the input sequence is then sampled. **Model Evaluation** A motif can be identified if the contribution scores higher than for random positions. To quantify how well a model captured a motif, we used an approach similar to [15]. Motif positions in the input sequence are labeled as positive, while random positions are labeled as negative. AUPRC values are then calculated based on those labels and the contribution scores similarly to prediction probabilities.

**Figure 5.**
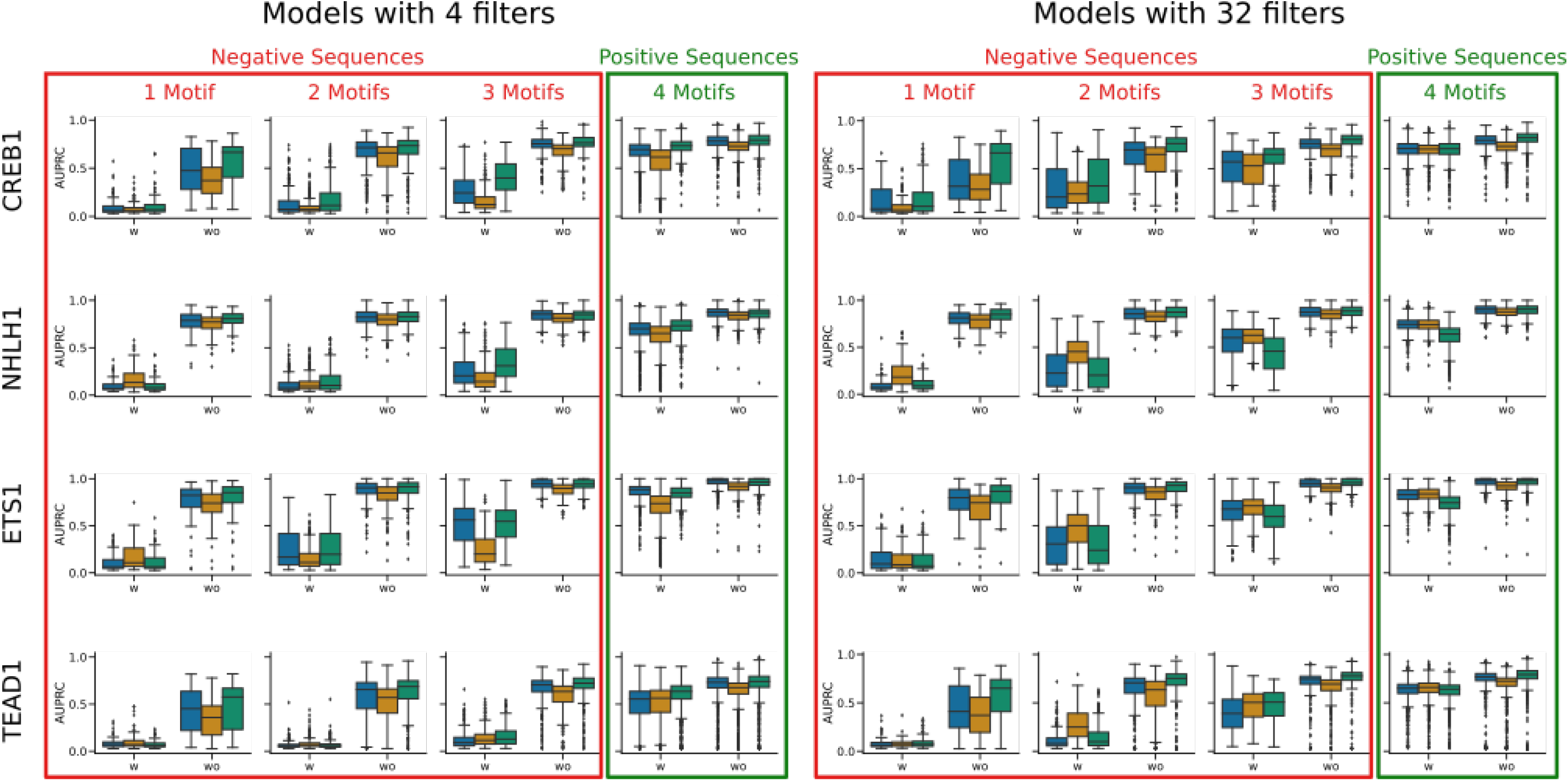
AUPRC scores for negative sequence experiment for distinct_1 motif set. AUPRC scores for the contribution scores of *M*_*w*_ are lower for all motifs compared to *M*_*wo*_.

**Figure 6.**
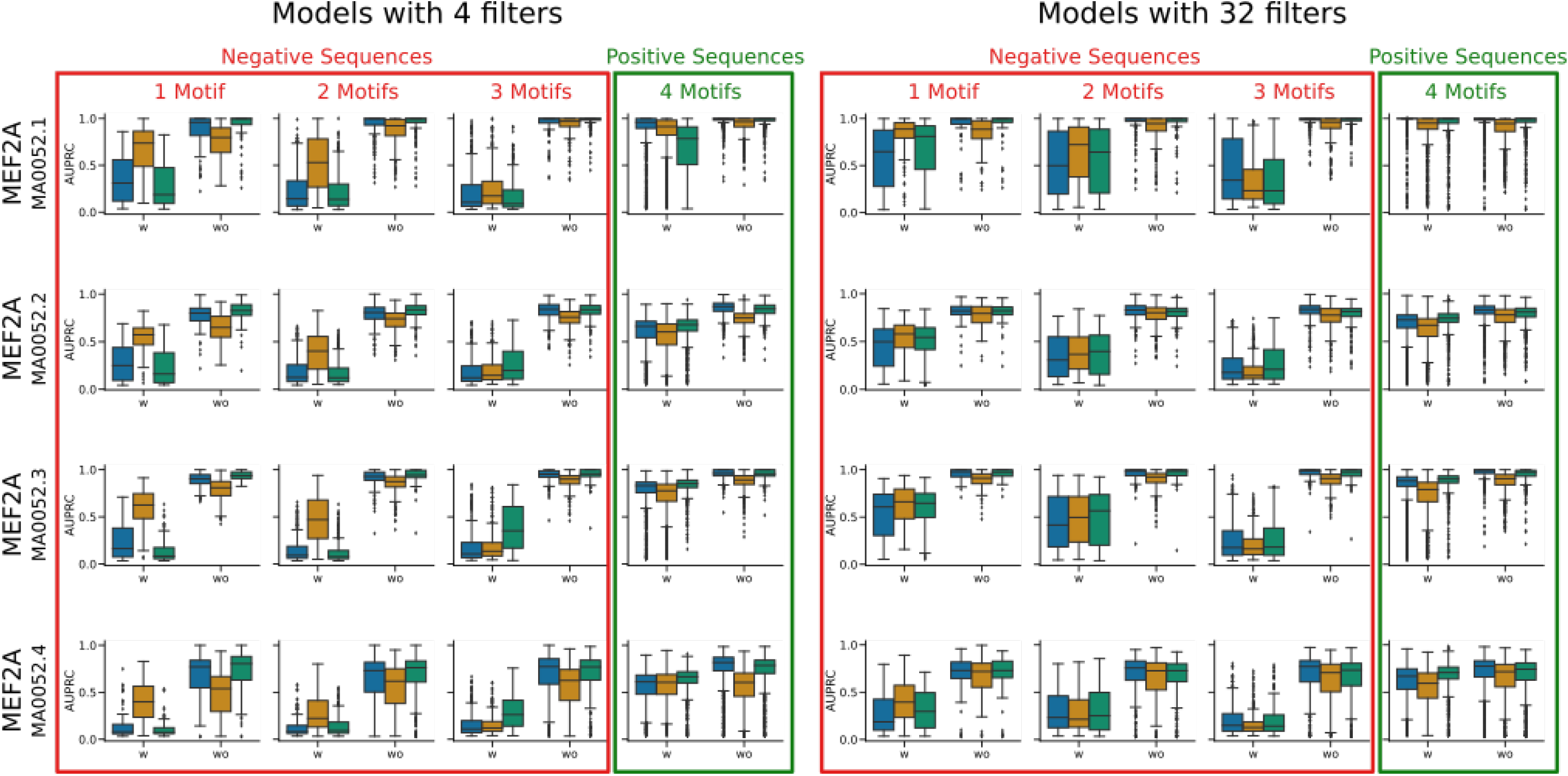
AUPRC scores for negative sequence experiment for MEF2A motif set. While the AUPRC values decrease with a lower number of motifs present in the case of heterologous motifs (see Supplement Fig. 5, AUPRC values are higher in the homologous case for the sequences containing 1 or 2 motifs compared to 3 since those sequences are closer to the decision boundary, especially for Feature Permutation.

**Figure 7.**
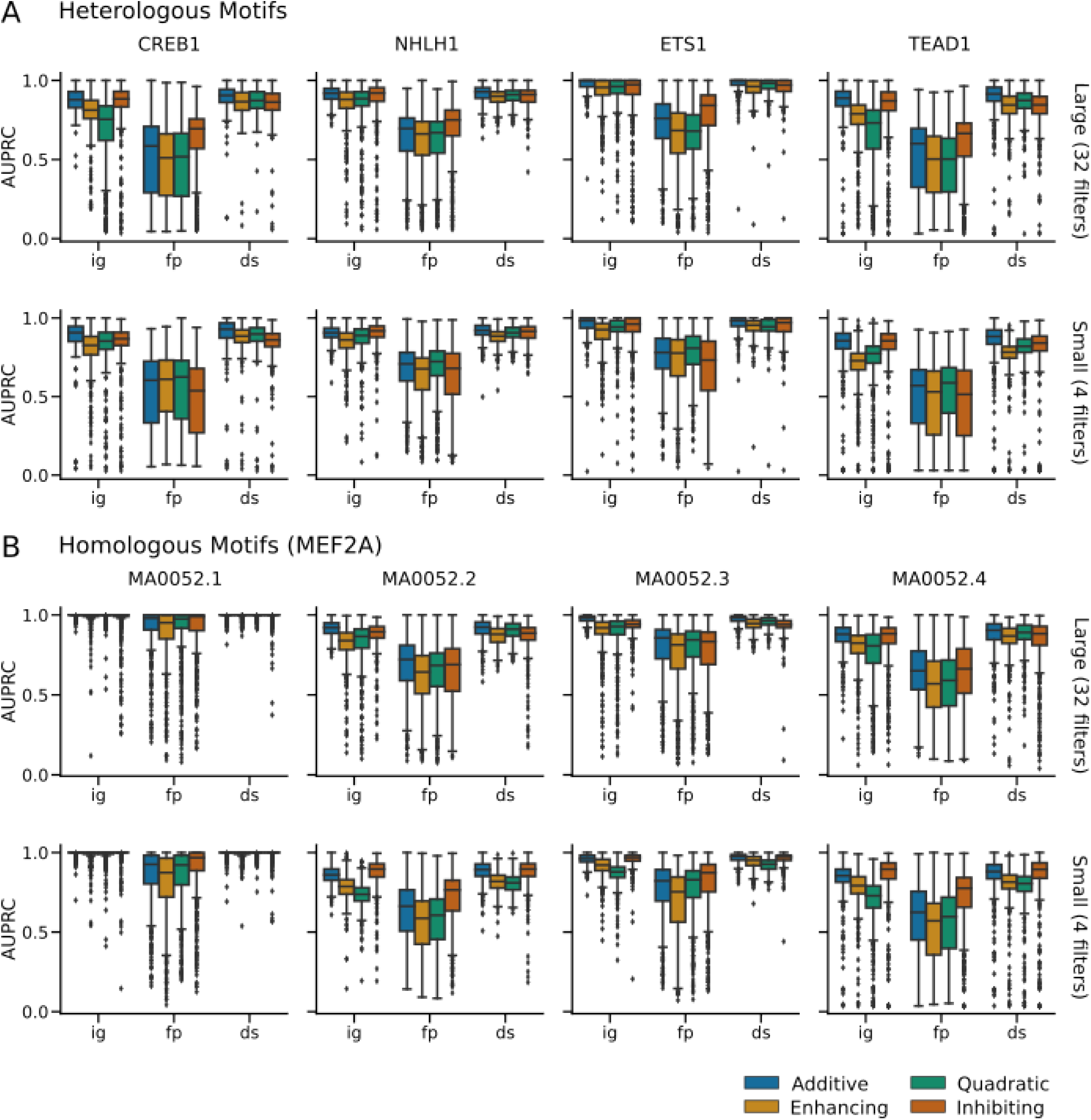
AUPRC for interaction effects for sequences containing all 4 motifs compared to additive effect (blue) for (A) an interactive motif set containing heterologous motifs and (B) a homologous motif set for MEF2A. DeepLIFT remains mostly consistent across the interaction effects whereas Integrated Gradients shows a decrease in motif detection for enhancing and quadratic interaction effects. Despite a similar interaction definition and model performance (see Table 6 and 7), differences between enhancing and inhibiting effects can be observed. There are also differences between the motifs in one set, even for the homologous data set which consists of similar motifs and therefore should yield similar contribution scores.

## Code availability

Code for the evaluation is available under www.gitlab.com/dacs-hpi/interpret-interaction upon publication.

## Author contributions statement

MSL designed the experiment. MSL, JMB, and BYR consulted on analytical decisions. MSL performed the analysis and wrote the manuscript with input from all co-authors. The authors read and approved the final manuscript.

## Appendix

